# Identification, Characterization and Mode of Action of Corynaridin, a Novel Linaridin from *Corynebacterium lactis*

**DOI:** 10.1101/2022.05.11.491181

**Authors:** Efthimia Pashou, Sebastian J. Reich, Dominik Weixler, Bernhard J. Eikmanns, Christian U. Riedel, Oliver Goldbeck

**Author notes:** Corresponding authors:, Address: Institute of Microbiology and Biotechnology, University of Ulm, Albert-Einstein-Allee 11, 89081 Ulm, Germany.

## Abstract

Genome analysis of *Corynebacterium lactis* revealed a bacteriocin gene cluster encoding a putative bacteriocin of the linaridin family of ribosomally synthesized and posttranslationally modified peptides (RIPPs). The locus harbors typical linaridin modification enzymes but lacks genes for a decarboxylase and methyltransferase, which is unusual for type B linaridins. Supernatants of *Corynebacterium lactis* RW3-42 showed antimicrobial activity against *Corynebacterium glutamicum*. Deletion of the precursor gene *crdA* clearly linked the antimicrobial activity of the producer strain to the identified gene cluster. Following purification, we observed potent activity of the peptide against Actinobacteria, mainly other members of the genus *Corynebacterium* including the pathogenic species *Corynebacterium striatum* and *Corynebacterium amycolatum*. Also, low activity against some Firmicutes was observed, but no activity against Gram-negative species. The peptide is resilient towards heat but sensitive to proteolytic degradation by trypsin and proteinase K. Notably, time-kill kinetics and experiments using live biosensors to monitor membrane integrity suggest bactericidal activity that does not involve formation of pores in the cytoplasmic membrane. As *Corynebacterium* species are ubiquitous in nature and include important commensal and pathogens of mammalian organisms, secretion of bacteriocins by species of this genus could be a hitherto neglected trait with high relevance for intra- and interspecies competition and infection.

**Importance:** Bacteriocins are antimicrobial peptides produced by bacteria to fend off competitors in ecological niches and are considered to be important factors influencing the composition of microbial communities. However, bacteriocin production by bacteria of the genus *Corynebacterium* has been a hitherto neglected trait, although its species are ubiquitous in nature and make up large parts of the microbiome of humans and animals. In this study, we describe and characterize a novel linaridin-family bacteriocin from *Corynebacterium lactis* and show its narrow spectrum activity, mainly against other actinobacteria. Moreover, we were able to extend the limited knowledge on linaridin bioactivity in general and for the first time describe the mode of action of such a bacteriocin. Interestingly, the peptide, which was named corynaridin, appears bactericidal, but without formation of pores in the bacterial membrane.

## 1 Introduction

Genome sequencing and bioinformatics have promoted the discovery of ribosomally synthesized, bioactive molecules over the past decades. The heterogeneous group of antimicrobial peptides produced by bacteria, so called bacteriocins, gained special interest because of their potential use as food preservatives and alternatives to antibiotics (1–5). However, the primary biological function of bacteriocins is to provide the producer with a selective advantage over target organisms in a complex and competitive ecological niche (6).

While bacteriocins have been extensively studied in lactic acid bacteria (LAB), comparably little knowledge is available about production of such compounds by other bacteria, e.g. Actinobacteria. Nevertheless, several studies suggest the widespread occurrence of bacteriocin gene clusters (BGCs) in non-LAB species, including the genus *Corynebacterium* (7–10). Species of this genus are wide-spread in nature, make up one of the largest groups of bacteria in the human and animal skin microbiome and are also present in food products including raw milk or cheese (11–13).

Besides toxicogenic *Corynebacterium* species, e.g. *Corynebacterium diphtheriae* and *Corynebacterium ulcerans*, many (non-diphtheritic) species of the genus have been described as commensals (12). Being a dominant bacterial group of the human skin microbiome, also non-diphtheritic corynebacteria are regularly found in infectious tissue. However, in most cases they are regarded rather a contamination from surrounding skin than the etiological agent of the infection itself (14). Nevertheless, improved methods to discriminate infection and colonization and an increasing number of reports suggest that some *Corynebacterium* species are important opportunistic pathogens for humans and animals (15). The clinical relevance of non-diphtheritic corynebacteria becomes even more apparent with increasing reports of multi-drug-resistant strains, mostly identified in nosocomial environments (16, 17). Species like *Corynebacterium striatum* and *Corynebacterium amycolatum* have been described to cause infections in elderly, immunocompromised patients and are associated with chronic wounds (18). Also, recent studies suggest that microbe-microbe interactions of *Corynebacterium* species with other commensals or pathogens like *Staphylococcus aureus* might influence the behavior and fitness of both species (19). Interestingly, only very few reports on bacteriocin production in corynebacteria exist (20, 21).

In general, bacteriocins of Gram-positive bacteria can be classified into small (<10 kDa) modified (class I) and unmodified (class II) as well as larger (>10 kDa) heat labile (class III) peptides/proteins (1). Class I bacteriocins usually contain posttranslational amino acid modifications such as dehydration, heterocycle formation, glycosylation, methylation, etc., that are often important for their biological activity (22). Thus, class I bacteriocins are also referred to as ribosomally synthesized and posttranslationally modified peptides (RiPPs) (1). Linaridins are a group of RiPPs with an overall linear structure, containing dehydrated amino acids such as dehydrobutyrine (23). So far, only five members of this family have been described in detail (cypemycin, grisemycin, legonaridin, mononaridin and salinipeptins) (23–29). Nevertheless, *in silico* analyses suggest that linaridin BGCs are widespread in nature and especially in Actinobacteria (7). In contrast to lanthipeptides, which also contain dehydrated amino acids (e.g. nisin), linaridin biosynthesis is considered to be essentially different from other RiPPs. Modification of the prototypic type A linaridin cypemycin includes dehydration of threonine residues, N-terminal methylation, C-terminal oxidative decarboxylation of cysteine and subsequent formation of a heterocyclic S-[(Z)-2-aminovinyl]-D-cysteine (AviCys) moiety (23, 24). N-terminal methylation was shown to be crucial for the activity of cypemycin (27). In contrast, type B linaridin gene clusters do not encode decarboxylases and thus lack C-terminal modification. Instead, genes for so far uncharacterized short-chain oxidoreductases have been identified, e.g. in the gene cluster for the biosynthesis of legonaridin (26).

While extensive studies have been carried out on the structure and chemistry behind their modifications, comparably little is known about the biological functions of linaridins. Antimicrobial activity of cypemycin as well as legonaridin appears to be limited to *Micrococcus luteus* (25, 26). Additionally, cypemycin possesses cytotoxic activity against mouse P388 leukemia cells (25, 28). The structurally similar salinipeptins, however, inhibit growth of a *Streptococcus pyogenes* strain but not of *M. luteus* (29). So far, for none of the hitherto described linaridins a receptor or mode of action has been proposed.

In this study, we describe the identification and partial characterization of a novel linaridin discovered in *Corynebacterium lactis* RW3-42, a strain isolated from raw cow’s milk (30).

## 2 Materials and Methods

### 2.1 Bacterial strains and cultivation conditions

**Table 1:** Bacterial strains and plasmids used in this study.

| Species | Strain | Source/Reference |
| --- | --- | --- |
| *Bacillus subtilis* | DSM 402 | DSMZ<sup>b</sup> |
| *Corynebacterium lactis* | RW3-42 | (30) |
| *Corynebacterium lactis* | RW3-42 ΔcrdA | this study |
| *Corynebacterium ammoniagenes* | DSM 20306 | DSMZ<sup>b</sup> |
| *Corynebacterium amycolatum* | DSM 6922 | DSMZ<sup>b</sup> |
| *Corynebacterium canis* | DSM 45402 | DSMZ<sup>b</sup> |
| *Corynebacterium casei* | DSM 44701 | DSMZ<sup>b</sup> |
| *Corynebacterium efficiens* | DSM 44549 | DSMZ<sup>b</sup> |
| *Corynebacterium glutamicum* | ATCC 13032 | ATCC<sup>a</sup> |
| *Corynebacterium lipophiloflavum* | DSM 44291 | DSMZ<sup>b</sup> |
| *Corynebacterium striatum* | DSM 20668 | DSMZ<sup>b</sup> |
| *Corynebacterium xerosis* | DSM 20743 | DSMZ<sup>b</sup> |
| *Cutibacterium acnes* | DSM 16379 | DSMZ<sup>b</sup> |
| *Escherichia coli* | MG1655 | (31) |
| *Lactiplantibacillus plantarum* | DSM 1055 | DSMZ<sup>b</sup> |
| *Lactococcus lactis* | IL1403 | (32) |
| *Listeria innocua* | LMG2785 | (33) |
| *Listeria monocytogenes* | EGD-e | (34) |
| *Micrococcus luteus* | DSM 20030 | (35) |
| *Pediococcus acidilactici* | 347 | (36) |
| *Pseudomonas fluorescens* | DSM 50090 | DSMZ<sup>b</sup> |
| *Staphylococcus aureus* | ATCC 29213 | ATCC<sup>a</sup> |
| *Staphylococcus epidermidis* | DSM 3269 | (37) |
| Plasmid | Characteristics |  |
| pk19mobsacB | Km<sup>r</sup>; mobilizable (oriT); oriV | (38) |
| pk19mobsacB_del-crdA | pK19-derivative with 750 bps flanking regions of the *crdA* gene of the *C. lactis* RW3-42 genome | this study |
| pPB-pHin2<sup>Cg</sup> | pPBEx2 derivative with codon optimized pHluorin2 gene under control of P<sub>tuf</sub>, Km<sup>r</sup>, pMB1 origin, pBL1 origin | (39) |
| pBAD33 | Cm<sup>r</sup>; pACYC184/p15A origin; *araC* | (40) |
| pBAD33_crd | pBAD33 derivative with *crd* gene cluster under control of P<sub>BAD</sub> | this study |

<sup>a</sup>: ATCC – American Type Culture Collection
<sup>b</sup>: DSMZ – Deutsche Sammlung von Mikroorganismen und Zellkulturen

### 2.2 Cultivation conditions

Bacterial strains used in this study (Table 1) were cultivated in GM17 medium (*L. lactis* IL1403), MRS medium (*L. plantarum* DSM1055, *P. acidilactici* 347) or BHI medium (all other) at 37°C (*E. coli* MG1655, *L. innocua* LMG2785, *L. monocytogenes* EGD-e, *S. aureus* ATCC 29213, *S. epidermidis* DSM 3269, *C. acnes* DSM 16379, *C. canis* DSM 45402, *C. efficiens* DSM 44569, *C. amycolatum* DSM 6922, *C. striatum* DSM 20668, *C. lipophiloflavum* DSM 44291) or 30°C (all other), respectively. Solidified medium was prepared by addition of 16 g agar per L to the medium. For growth characterization and production, *C. lactis* strains were cultivated in *Corynebacterium lactis* medium (CLI) (21 g/L MOPS, 1 g/L K_2_HPO_4_, 1 g/L KH_2_PO_4_, 16 g/L tryptone, 10 g/L yeast extract, 0.25 g/L MgSO_4_, 0.01 g/L CaCl_2_, 0.2 mg/L biotin, pH adjusted to 7). For time kill kinetics, 2xTY medium (16 g/L tryptone, 10 g/L yeast extract, 5 g/L NaCl) was used for the cultivation of *C. glutamicum* ATCC 13032. Cultivation of strains carrying plasmids additionally contained kanamycin (25 µg/mL) or chloramphenicol (15 µg/mL). Growth was monitored photometrically by measuring the optical density at 600 nm (OD_600_) at the indicated time points.

### 2.3 Molecular biology procedures

Construction of pk19mobsacB_del-crdA and pBAD33_crd was carried out with standard reagents and according to protocols of the manufacturers. Oligonucleotides are listed in Table S1 and were obtained from Eurofins Genomics (Ebersberg, Germany). Polymerase chain reaction (PCR) was performed in a C100 thermocycler (Bio-Rad Laboratories, Munich, Germany) using Q5 high fidelity polymerase (New England Biolabs, Ipswich, USA) and nucleotides from Bio-Budget (Krefeld, Germany). The up- and downstream region of the *crdA* gene and the *crd* gene cluster were amplified using oligonucleotides creating overlapping ends for Gibson Assembly®. The empty vector pK19mobsacB was linearized by restriction endonucleases *EcoR*I and *Sal*I. The empty vector pBAD33 was linearized by restriction endonucleases *Kpn*I and *Sal*I. The final plasmids were verified by sequencing (Eurofins Genomics, Ebersberg, Germany). All plasmids and their relevant characteristics are listed in Table 1. For transformation, *C. lactis* was rendered electrocompetent and transformed as described previously for *C. glutamicum* (41).

### 2.4 In silico analyses

Prediction of bacteriocin gene clusters (BGCs) in the genome of *C. lactis* RW2-5 was carried out using BAGEL4 (42). Subsequently, BLASTP analysis (43) was performed using the deduced proteins with standard parameters (BLOSUM62; gap existence costs: 11; gap extension costs: 1) and assigned to putative functions based on sequence similarities with published linaridin biosynthesis genes (23). The BGC of *C. lactis* RW3-42 was PCR-amplified and cloned into pBAD33, followed by sequencing (Eurofins Genomics, Ebersberg, Germany). Resulting sequences were then aligned to the BGC of *C. lactis* RW2-5 and the legonaridin biosynthesis genes. Sequence alignments were carried out using ClustalW (44) and visualized using Jalview (45).

### 2.5 Purification of corynaridin

For purification of corynaridin, protein of supernatants (1 L) collected after 24 h of cultivation of *C. lactis* RW3-42 in CLI containing 1 % (w/v) glucose were precipitated using ammonium sulfate (50 % (w/v) saturation) at 4 °C overnight (16 h). The precipitate was collected by centrifugation (60 min, 10,000 g, 4 °C), re-suspended in 50 mL H_2_O, and pH was adjusted to 4.0 using 2 M HCl. An additional centrifugation step (10 min, 10,000 g, 4 °C) was performed to remove insoluble particles. All following chromatographic steps were carried out with an ÄKTA pure chromatography system (Cytiva). The solution containing the peptide was applied to a HiPrep SP FF 16/10 column (GE Healthcare Life Sciences) equilibrated with 20 mM sodium phosphate buffer at pH 3.9. Unbound proteins were washed out by 5 column volumes (CVs) of 20 mM sodium phosphate buffer at pH 6.9. Remaining bound peptides/proteins were then eluted in with 5 CVs of 20 mM sodium phosphate buffer at pH 6.9 with 2 M NaCl. The eluate fractions containing the bacteriocin were identified by activity assays (see below) and directly applied to reversed phase chromatography (RPC) using a 1 mL Resource RPC column (Cytiva) or stored at -20 °C until further use. To remove weakly bound proteins, a washing step was carried out with 5 CVs of 2 % acetonitrile in H_2_O+ 0.065 % TFA. Elution was performed with an initial step to 15 % acetonitrile for 5 CVs. followed by a linear gradient up to 80 % acetonitrile. Fractions with antimicrobial activity were identified by activity assays (see below), dried using a vacuum concentrator at 60°C (Eppendorf, Hamburg, Germany) and re-suspended in HPLC-grade H_2_O. Protein concentrations of the purification fractions were estimated using the Pierce™ BCA Protein Assay Kit (Thermo Fisher Scientific) according to the manufacturers protocol.

### 2.6 Radial streak

Bioprospecting for antimicrobial activity was performed using a modified cross-streak method (46). Briefly, *C. lactis* RW3-42 was inoculated from an overnight (o/N) culture as a single streak with an inoculation loop in the center of an BHI media agar plate and incubated aerobically for at least 3 days at 30°C. Indicator bacteria (Table 1) were cultivated overnight in 5 mL BHI media and streaked in a line from the border of the plate towards *C. lactis*. The plates were then incubated for 24 to 48 h at 30 to 37°C depending on the indicator bacteria. In case of *Cutibacterium acnes* incubation was carried out in an anaerobic jar (Merck KGaA, Darmstadt, Germany) containing AnaeroGen™ anaerobic incubation system (Thermo Fisher Scientific).

### 2.7 Determination of antimicrobial activity

Antimicrobial activity was determined using a spot-on-lawn assay. Overnight cultures of the respective strains were inoculated with an OD_600_ of 0.01 into hand-warm agar medium (16 g/L agar) and poured into sterile petri dishes. After solidification, surfaces of agar plates were air-dried at RT under a sterile hood. Supernatants and purification fractions were serially diluted and spotted onto agar plates. Plates were incubated under the preferred conditions of the embedded bacteria until growth was visible. Volumetric bacteriocin activity (bacteriocin units per mL; BU/mL) was determined by dividing the last dilution factor resulting in a visible zone of inhibition by the volume spotted.

### 2.8 Time-kill kinetics

Fresh o/N cultures of *C. glutamicum* ATCC 13032 were used to inoculate 5 mL 2xTY in glass tubes with a starting OD_600_ of 0.5, i.e. approximately 10^7^ CFU/mL. Samples of interest were added at the indicated concentrations prior to inoculation of the medium. Cultures were then incubated on a rotary shaker at 130 rpm for 24 h at 30 °C. Samples were collected at the indicated time points, diluted (10^−1^ – 10^−8^) and plated on 2xTY agar. Colony forming units per mL (CFU/mL) were determined after 24 - 48h of incubation of the plates at 30°C by counting the colonies for the respective dilution.

### 2.9 pHluorin assay

For detection of membrane damage, a pHluorin assay was conducted as described earlier (39). In particular, a 5 mL BHI overnight culture containing kanamycin (50 µg/mL) of the sensor strain *C. glutamicum*/pPB-pHin2^*Cg*^ was harvested by centrifugation and resuspended to an OD_600_ of 3 in Listeria minimal buffer (LMB; 100 mM MOPS, 4.82 mM KH_2_PO_4_, 11.52 mM Na_2_HPO_4_, 1.7 mM MgSO4, 0.6 g/L (NH_4_)_2_SO_4_, 55 mM glucose, pH 6.2). Serial 2-fold dilutions of samples were prepared in black 96 well microtiter plates (Sarsted, Nümbrecht, DE) with a final volume of 100 µL in each well. Subsequently, 100 µL of the sensor strain suspension was added and the plate was incubated at room temperature in the dark for 30 min. Then, pHluorin2 fluorescence was measured at 520 nm with excitation at the distinct maxima at 400 and 480 nm using an infinite M200 plate reader (Tecan, Männedorf, Swiss).

### 2.10 Fluorescence microscopy

A fresh culture of *C. glutamicum* ATCC 13032 was washed once in PBS and bacteria were resuspended in saline (0.9 % NaCl, w/v) at OD_600_ = 1. 87.5 µl of the cell suspension were mixed with nisin, cetyltrimethylammonium bromide (CTAB), H_2_O or RPC fraction containing corynaridin to the indicated concentrations and incubated for 10 - 30 min in the dark. Then, the bacteria were stained using 12.5 µl propidium iodide (25 µg/ml; Invitrogen, Darmstadt, Germany) and again incubated for 15 min in the dark. Samples were imaged using a Axio Observer Z1 (Zeiss, Oberkochen, Germany) in bright field and fluorescence mode with a filter set for propidium iodide (excitation at 575–625 nm; emission at 660–710 nm). Images were acquired with a 63× objective and analyzed using the Zen software (Version 2.3 SP1; Zeiss).

## 3 Results

### 3.1 In silico analysis of a bacteriocin gene cluster in C. lactis

*In silico* analyses using the web-based tool BAGEL4 revealed several, yet undescribed bacteriocin gene clusters (BGC) in the genus *Corynebacterium* (8). In this study, we closely examined one of the predicted BGCs in the genome of *C. lactis* RW2-5 isolated from raw cow’s milk and found an identical cluster in the strain *C. lactis* RW3-42 upon sequencing of the locus (30). The identified BGC consists of six genes, including a gene for a conserved LinL-protein homolog (here named *crdL*) that was so far only associated with linaridin biosynthesis and is typically used to identify corresponding gene clusters (Figure 1) (27). Furthermore, a hypothetical peptide precursor gene (here named *crdA*) was identified which encodes a peptide of 69 amino acids with a conserved hexapeptide cleavage motif PxxxTP at positions 29 – 34. The peptide also harbors a high number of threonine residues (seven in total) in its C-terminal part, which are often post-translationally modified in RiPPs as shown for cypemycin or nisin (Figure 1) (22).

**Figure 1:**
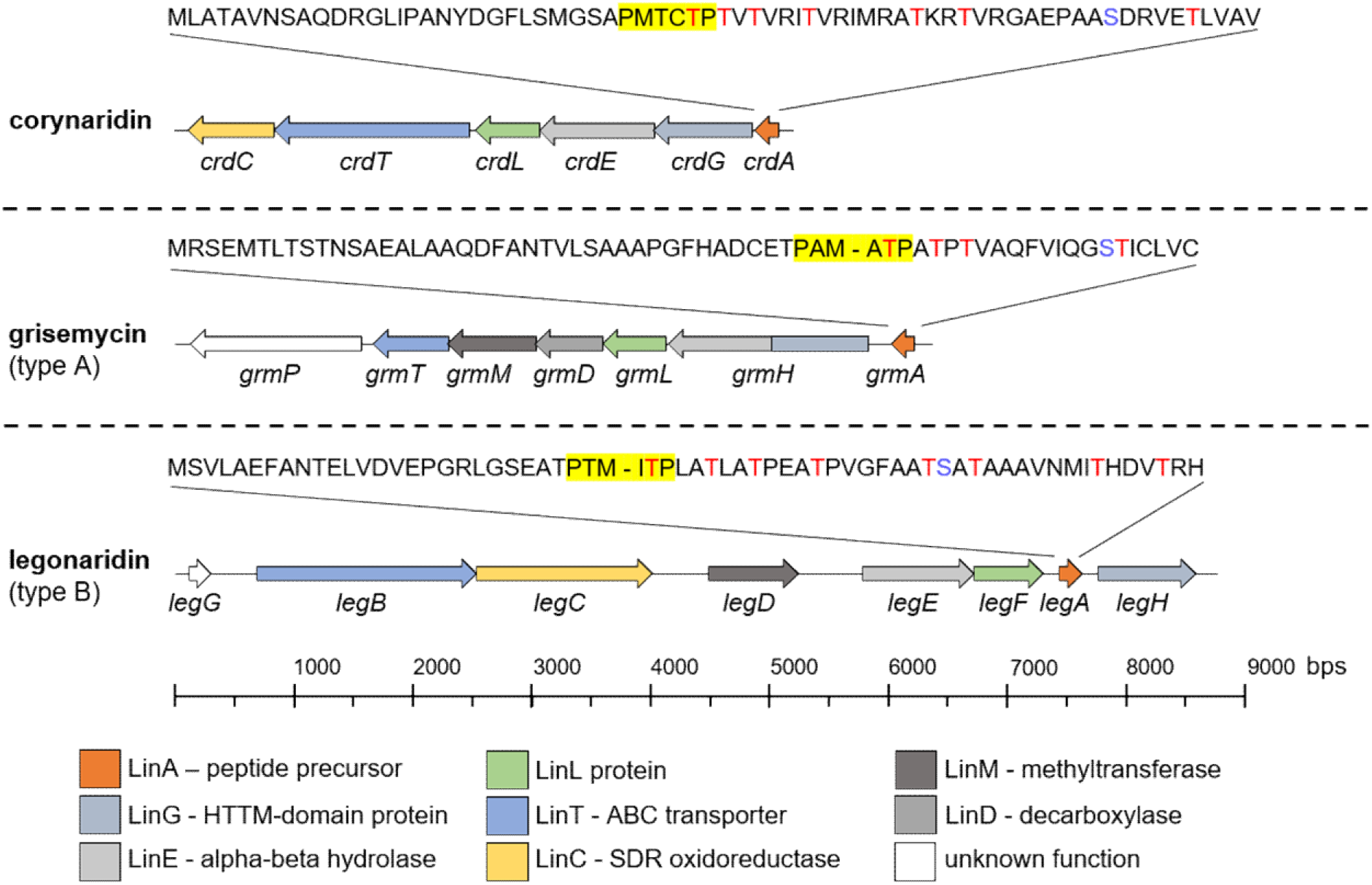
Comparison of the BGC of *C. lactis* RW3-42 with BGCs for other bacteriocins of the linaridin family. Genetic organization of the corynaridin, grisemycin and legonaridin BGCs as predicted by BAGEL4 and BLASTP analyses. The amino acid sequence of the prepeptides are displayed above the gene cluster. Threonine residues are displayed in red letters. The hexapeptide cleavage site PxxxTP in the prepeptides is highlighted in yellow. Predicted functions of modification enzymes and their corresponding genes are indicated by a color scheme according to the legend at the bottom of the figure.

The deduced amino acid sequences of the other proteins encoded in the BGC were further analyzed using BLASTP and checked for homologies to enzymes involved in biosynthesis of described members of the linaridin family. Similar to other linaridin BGCs, the *crd* locus contains no gene for a protease involved in removal of the leader peptide but genes for an N-terminal horizontally transferred transmembrane helix domain protein (HTTM-domain protein, *crdG*) and an alpha-beta-fold hydrolase (*crdE*) are present (Figure 1). These putative modification enzymes are assumed to participate in the maturation of linaridins although their exact mechanism remains unclear (23). Interestingly, no decarboxylase gene but a putative short-chain oxidoreductase (SDR, *crdC*) typical for biosynthetic operons of type B linaridins is part of the BGC. This observation and the absence of a C-terminal cysteine residue in the peptide precursor suggest that the BGC of *C. lactis* encodes a type B linaridin. Notably, we were also unable to identify a gene for a methyltransferase involved in N-terminal methylation, which was described for e.g. legonaridin (23, 26). The ABC-transporter (*crdT*) encoded in the BGC is supposedly involved in transport and/or processing of the peptide or immunity of the host (48). Sequence alignments of the enzymes of the *C. lactis* BGC with those of the legonaridin biosynthesis of *Streptomyces* sp. CT34 revealed only low overall homologies (22 - 32 %; Supplementary Table S2 and Figure S1 – S6). Overall, our *in silico* analyses suggest that the BGC of *C. lactis* RW3-42 encodes for a novel type B linaridin lacking N-terminal methylation, for which we propose the designation “corynaridin”.

We furthermore analyzed whether the corynaridin precursor exists in other BGCs of *Corynebacterium* species and found a highly similar gene cluster in the genome of *C. striatum* 1329_CAUR, isolated from the wound of an intensive care unit patient (Figure S7) (49). The respective peptide precursor shows 84 % identity to corynaridin with a conserved PxxxTP hexapeptide motif and seven putatively modified threonine residues at its N-terminus. Interestingly, a previous, extensive bioinformatic analysis of the NCBI genome database identified 561 linaridin BGCs mainly in Actinobacteria, including the ones in *C. lactis* and *C. striatum* plus four other in *Corynebacterium* spp. (7). Closer examination of these BGCs shows that also in these genomes typical linaridin biosynthesis genes are present, but the overall genetic architecture and precursor peptide sequences are significantly different (Figure S8 & 9).

### 3.2 Growth, antimicrobial activity and kinetic of secretion of corynaridin

Since bacteriocins are often most effective against closely related species, we first tested if *C. lactis* RW3-42 is able to inhibit growth of different *Corynebacterium* species. Analysis of its antimicrobial capacity against *C. glutamicum* ATCC 13032 revealed a clear zone of inhibition in cross streak and spot-on-lawn assays using supernatants of *C. lactis* RW3-42 (Figure 2a). Also, other tested *Corynebacterium* species, *M. luteus* DSM 20030 and *Pediococcus acidilactici* 347 were inhibited in growth (Table 2 and Table S3). By contrast, no activity of *C. lactis* RW3-42 was detected against *Cutibacterium acnes* DSM 16379, the other tested Firmicutes or Gram-negative bacteria like *Escherichia coli* MG1655 and *Pseudomonas fluorescens* DSM 50090 in cross streak assays (Table 2 and Table S3). To verify that the antimicrobial activity is related to the predicted BGC (Figure 1), we generated the mutant strain *C. lactis* Δ*crdA* with a clean, markerless deletion of *crdA*, encoding the corynaridin precursor. The mutant did not show antimicrobial activity against *C. glutamicum* ATCC 13032 in cross streak or spot-on-lawn assays confirming that the antimicrobial activity of *C. lactis* RW3-42 is related to *crdA* (Figure 2a).

**Table 2:** Inhibitory spectrum of *C. lactis* RW3-42 and purified corynaridin assessed with cross-streak and spot-on-lawn assays.

| Strain | Phylum | Activity<sup>a</sup> |  |
| --- | --- | --- | --- |
|  |  | Cross streak | RPC fraction<sup>b</sup> |
| *Corynebacterium ammoniagenes* DSM 20306 | Actinobacteria | + | + |
| *Corynebacterium amycolatum* DSM 6922 | Actinobacteria | + | + |
| *Corynebacterium canis* DSM 45402 | Actinobacteria | (+) | n.d. |
| *Corynebacterium casei* DSM 44701 | Actinobacteria | + | + |
| *Corynebacterium efficiens* DSM 44549 | Actinobacteria | + | + |
| *Corynebacterium glutamicum* ATCC 13032 | Actinobacteria | + | + |
| *Corynebacterium lipophiloflavum* DSM 44291 | Actinobacteria | + | + |
| *Corynebacterium striatum* DSM 20668 | Actinobacteria | + | + |
| *Corynebacterium xerosis* DSM 20743 | Actinobacteria | + | + |
| *Cutibacterium acnes* DSM 16379 | Actinobacteria | - | (+) |
| *Micrococcus luteus* DSM 20030 | Actinobacteria | + | + |
| *Bacillus subtilis* DSM 402 | Firmicutes | - | - |
| *Lactobacillus plantarum* DSM 1055 | Firmicutes | - | - |
| *Lactococcus lactis* IL1403 | Firmicutes | - | + |
| *Listeria innocua* LMG2785 | Firmicutes | - | (+) |
| *Listeria monocytogenes* EGD-e | Firmicutes | - | - |
| *Pediococcus acidilactici* 347 | Firmicutes | + | - |
| *Staphylococcus aureus* ATCC 29213 | Firmicutes | - | - |
| *Staphylococcus epidermidis* DSM 3269 | Firmicutes | - | - |
| *Pseudomonas fluorescens* DSM 50090 | Gammaproteobacteria | - | - |
| *Escherichia coli* K12 MG1655 | Proteobacteria | - | - |

<sup>a</sup> + = zone of inhibition, (+) = diffuse zone of inhibition, - = no zone of inhibition, n.d. = not determined
<sup>b</sup>10 µL RPC fraction

**Figure 2:**
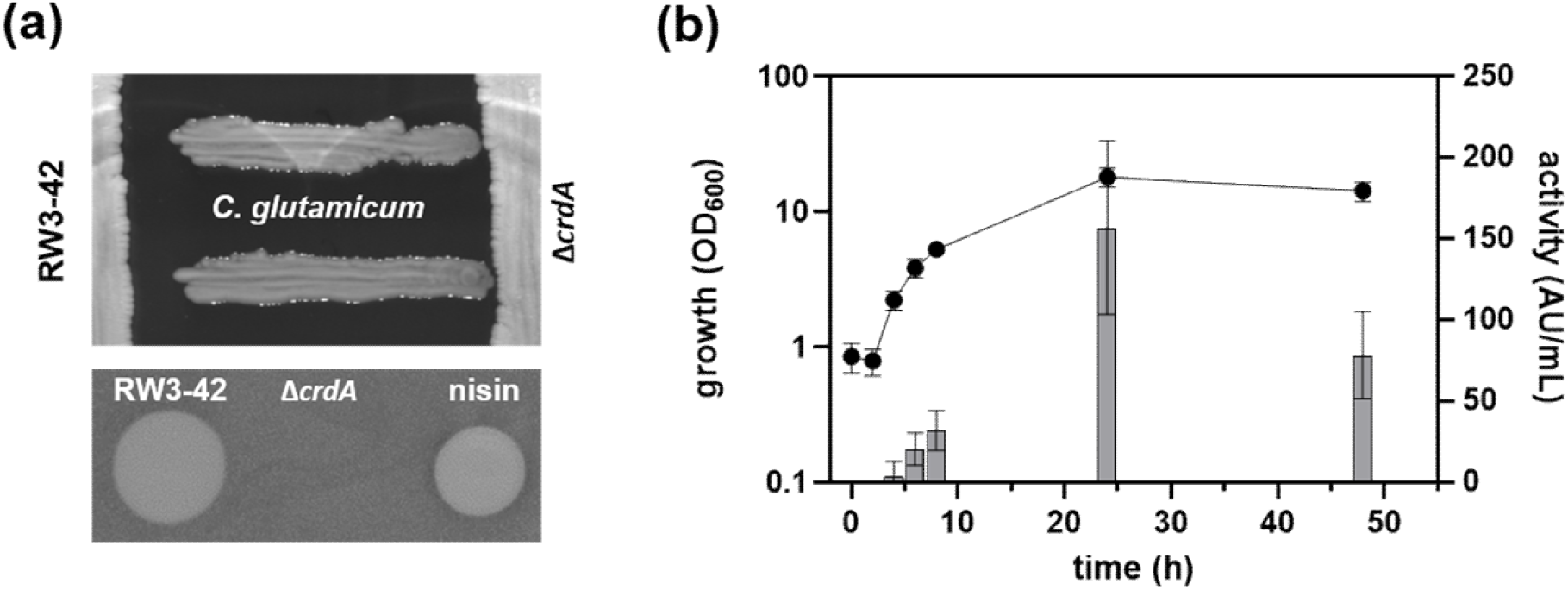
Analysis of the antimicrobial activity produced by *C. lactis* RW3-42. (a) Cross streak assay (upper panel) and spot-on-lawn assays (lower panel) indicating the secretion of an antimicrobial substance by *C. lactis* RW3-42 compared to the deletion strain *C. lactis* Δ*crdA* using *C. glutamicum* ATCC 13032 as indicator. 10 μL of a nisin standard (250 μg/mL) was used as a positive control in the spot-on-lawn assay. (b) Growth (OD_600_; left y-axis; black dots) and kinetic of antimicrobial activity (BU/mL; right y-axis; grey columns) of *C. lactis* RW3-42 in CLI medium containing 1 % (w/v) glucose. Antimicrobial activity was measured with *C. glutamicum* ATCC 13032 as indicator.

Growth of *C. lactis* RW3-42 and kinetics of production of the antimicrobial activity in shake flask experiments indicated that the antimicrobial compound is secreted mainly during exponential growth phase (Figure 2b). The highest biomass of the strain was observed after 24 h (OD_600_ = 18 ± 3). Minor antimicrobial activity against *C. glutamicum* ATCC 13032 was first observed after 4 h and peaked at a maximum of 157 ± 53 BU/mL after 24 h. After 48 h, a reduction of antimicrobial activity occurred, possibly due to degradation or adsorption of the peptide to biomass.

### 3.3 Purification of corynaridin from supernatants of C. lactis RW3-42

We next sought to purify the secreted antimicrobial compound for further characterization. Supernatants of 1 L cultivations of *C. lactis* RW3-42 were harvested and proteins were precipitated using ammonium sulfate. The precipitate was resuspended in HPLC grade water, pH adjusted to 4 and used for cation exchange chromatography (CIEX; Figure 3a). A single peak was observed following onset of elution with high-salt buffer and the corresponding fractions exhibited activity against *C. glutamicum* ATCC 13032 in spot-on-lawn assays. Peak fractions were pooled for further purification via reversed phase chromatography (RPC) using Acetonitrile-H_2_O-TFA as mobile phase (Figure 3b). A combined step and linear gradient was applied and yielded several peaks of which only one (at approx. 56 % acetonitrile) showed activity against *C. glutamicum* ATCC 13032 in spot-on-lawn assays. After removal of acetonitrile and resuspension in HPLC-grade H_2_O, a preparation with high activity (32,000 BU/mL) was obtained.

**Figure 3:**
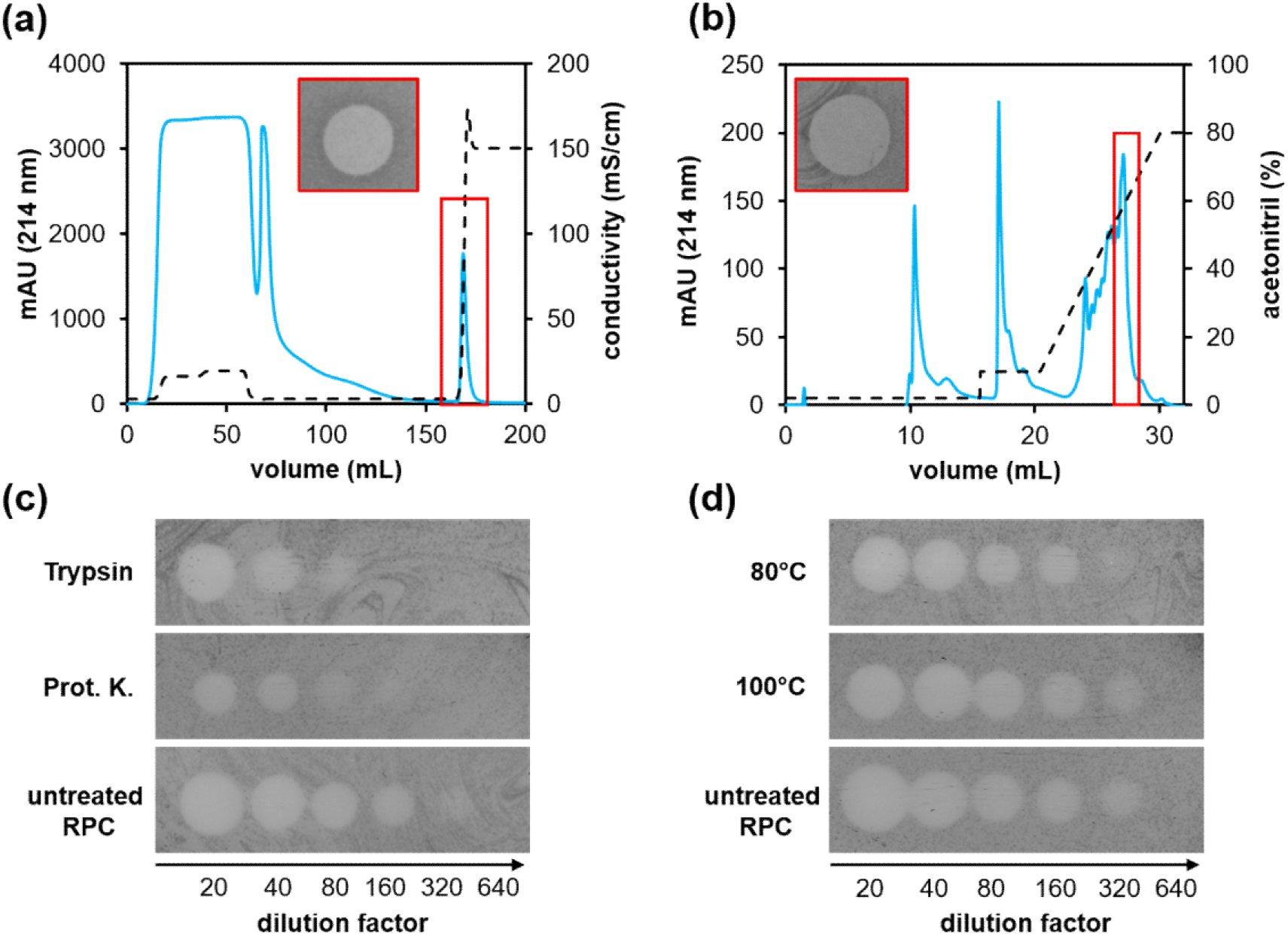
Purification of corynaridin from *C. lactis* RW3-42 supernatants. Supernatant proteins were precipitated by 50 % (w/v) ammonium sulfate and analyzed by (a) cation-exchange chromatography and (b) reversed-phase chromatography. Absorbance at 214 nm is displayed as blue line, conductivity and acetonitrile concentration are displayed as dashed line, respectively. Purified corynaridin (i.e. pooled fractions of the indicated peak of revers-phase chromatography) was analyzed for stability against (c) protease and (d) heat treatment. One representative experiment is shown for each purification step.

To assess the physicochemical properties of the purified compound, it was tested for resistance to protease and heat treatment (Figure 3c and d). While a reduction of activity of the RPC fraction to 8,000 BU/mL (i.e. about 4-fold) was observed after incubation with trypsin or proteinase K, heat treatment at 80 °C and 100 °C for 10 min had no effect. This indicates that the purified antimicrobial compound is a heat-stable peptide.

### 3.4 Inhibitory spectrum of purified corynaridin

Purified corynaridin was active against all tested *Corynebacterium* species (Table 2, Table S3) except *C. canis* DSM 45402 that could not be properly analyzed with the spot on lawn method. Notably, besides non-pathogenic, environmental and commensal bacteria including *C. glutamicum* ATCC 13032 (Figure 4a), also emerging multi-resistant pathogens like *C. striatum* DSM 20668 (Figure 4b) and *C. amycolatum* DSM 6922 (Figure 4c) were inhibited by RPC-purified corynaridin. In case of *P. acidilactici* 347, assays with purified peptide contradicted the cross-streak results as it did not inhibit growth of the strain (Table 2). *L. lactis* IL1403 was the only firmicute tested that was effectively inhibited by corynaridin. For *L. innocua* LMG2785 and the actinobacterium *C. acnes* DSM 16379 low levels of inhibition were achieved only at high concentrations (> 1000 µg/mL) of the RPC fraction.

**Figure 4:**
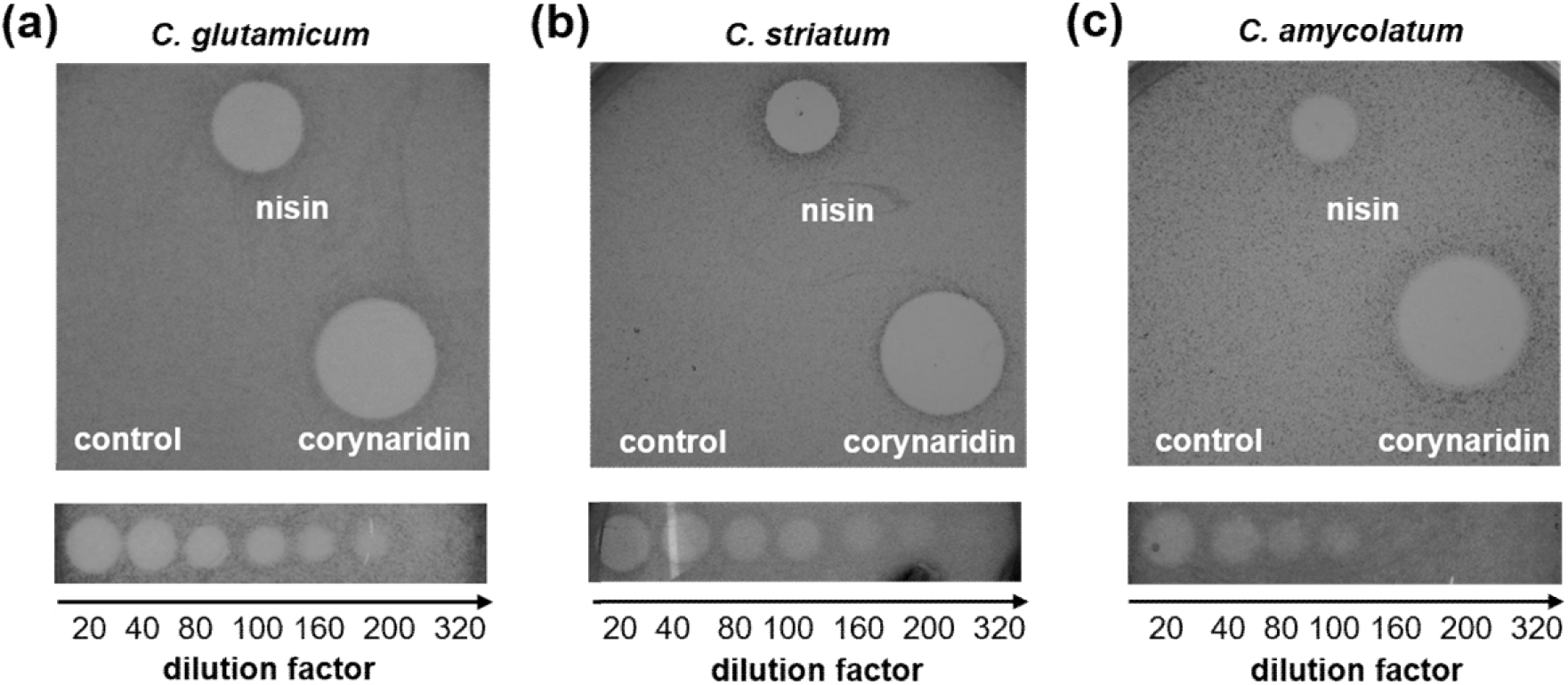
Antimicrobial activity of purified corynaridin against pathogenic corynebacteria. RPC fraction was used undiluted and in a dilution series against (a) *C. glutamicum* ATCC 13032, (b) *C. striatum* DSM 20668 and (c) *C. amycolatum* DSM 6922. For each spot, 10 μL sample were used. A nisin standard (250 μg/mL) and HPLC-grade H_2_O (control) were used as positive and negative controls, respectively.

For all other tested Gram-positive bacteria of the phylum Firmicutes, including *L. monocytogenes* EGD-e or *S. aureus* ATCC 29213, or Gram-negatives like *E. coli* MG1655 or *P. fluorescence* DSM 50090, cross-streak and spot-on-lawn assays both equally indicated that these organisms were not inhibited by corynaridin. Collectively, this suggests that corynaridin has a rather narrow spectrum of target organisms (Table 2).

### 3.5 Mode of action of corynaridin

To elucidate the mode of action of corynaridin, we employed time-kill assays with *C. glutamicum* ATCC 13032 as indicator. As controls for bacteriostatic and bactericidal compounds, we used the antibiotic chloramphenicol and the pore-forming class I bacteriocin nisin, respectively (Figure 5a). Addition of the vehicle H_2_O to *C. glutamicum* cultures had no effect on cell viability and CFU/mL steadily increased over the course of the experiment. By contrast, cultures treated with 1.25 µg/mL nisin showed a > 4-log reduced CFU/mL after 2 h cultivation as a consequence of the bactericidal effect of the peptide (Figure 5a). Addition of 6.5 µg/mL (bacteriostatic) chloramphenicol resulted in stable CFU/mL levels throughout the experiment. Corynaridin-containing RPC-fractions at 400 BU/mL and 1060 BU/mL decreased the CFU/mL of the indicator after 2 h by four orders of magnitude indicating bactericidal activity of the peptide. While cultures treated with the high concentration remained at low CFU/mL levels, those treated with the lower concentrations showed slightly increased CFU/mL after 24 h similar to nisin-treated samples indicating a bactericidal mode of action of corynaridin.

**Figure 5:**
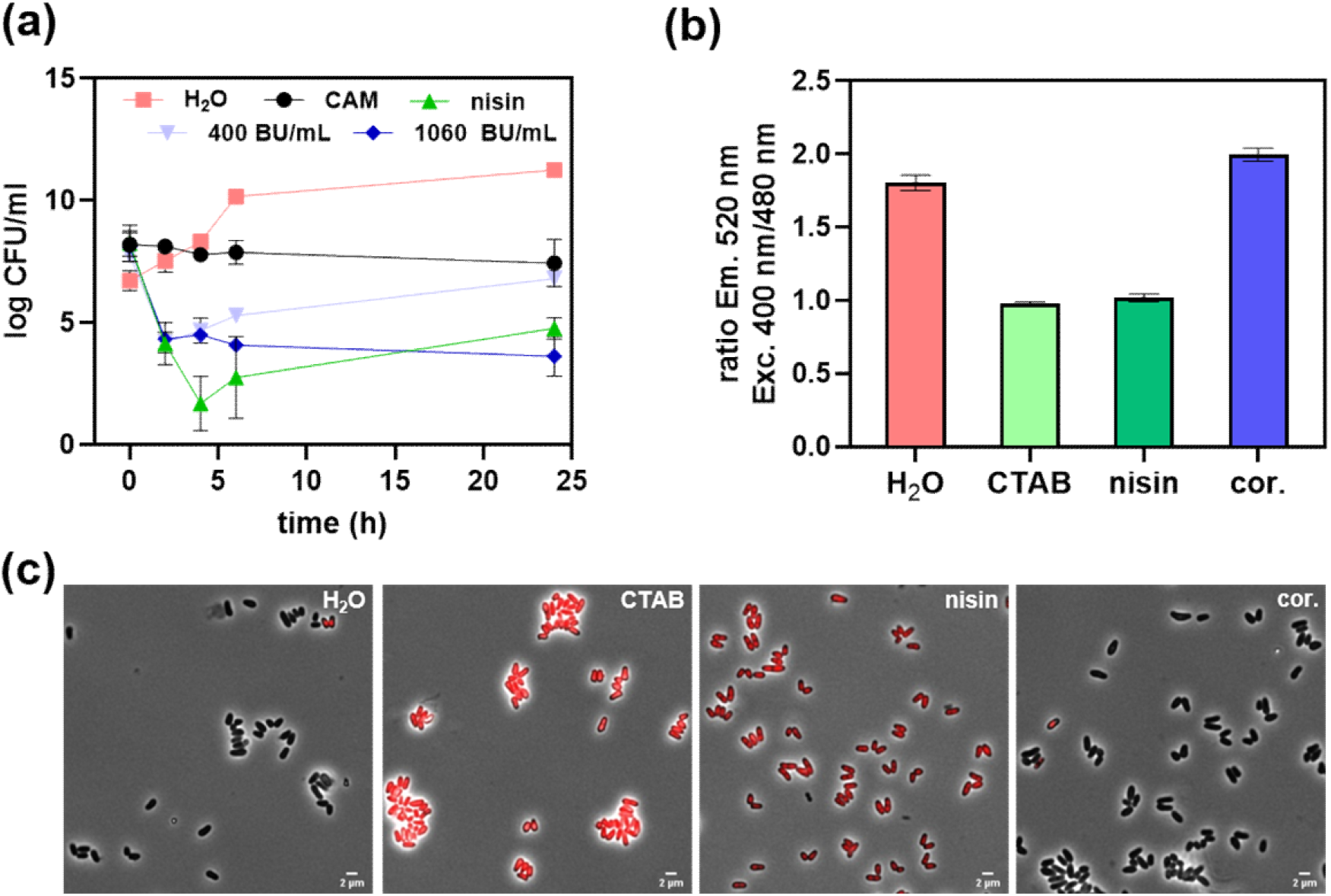
Mode of action of corynaridin. (a) Time-kill assays with *C. glutamicum* ATCC 13032 as indicator, 1.25 μg/mL nisin, 6.5 μg/mL chloramphenicol and H_2_O as controls and corynaridin (400 and 1060 BU/mL RPC fraction). (b) Fluorescence intensity ratios at a 400 and 480 nm excitation for H_2_O (neg. ctrl.), CTAB, nisin and corynaridin (cor.) added to *C. glutamicum*/pPB-pHin2^*Cg*^ cells for monitoring of membrane damage. (c) Fluorescence microscopic pictures (63x) of propidium iodide-stained *C. glutamicum* ATCC 13032 cells after treatment with H_2_O, 0.05 % CTAB, 1.25 μg/mL nisin, 4000 BU/mL RPC fraction containing corynaridin (cor).

As bactericidal activity of bacteriocins is often related to the formation of pores in the bacterial membrane, we tested whether corynaridin causes membrane damage using a fluorescence-based whole-cell biosensor assay with *C. glutamicum* ATCC 13032/pPB-pHin^Cg^ (39). The strain harbors a plasmid for expression of pHluorin2, a fluorescent protein with a pH-dependent, bimodal excitation spectrum which is highly suitable to determine bacteriocin-driven pore formation (47). As expected, the detergent CTAB and pore-forming nisin led to reduced fluorescence ratios for *C. glutamicum* ATCC 13032/pPB-pHin^Cg^ suggesting membrane damage (Figure 5b). By contrast, addition of the RPC-purified corynaridin preparation at concentrations up to 10.000 BU/mL did not result in a change of fluorescence ratios. These results were further supported by fluorescence microscopy of propidium iodide-stained bacteria, which provided no evidence for compromised integrity of the bacteria (Figure 5c). Thus, our data strongly suggests that unlike nisin and other bactericidal bacteriocins, corynaridin does not act by damaging the membrane of target cells under the tested conditions but by an as yet unknown bactericidal mode of action.

## 4 Discussion

In this study, we identified a putative linaridin BGC in the genome of *C. lactis* RW3-42, which was isolated from raw cow’s milk (30). Deletion of the peptide precursor gene *crdA* in the genome of *C. lactis* RW3-42 completely abolished antimicrobial activity of the strain and thus confirmed the identified BGC as locus for the biosynthesis of an antimicrobial compound, which we designated corynaridin. Further *in silico* analyses revealed several adjacent genes for putative modification enzymes that are typically associated with the linaridin family of RiPPs (23). Absence of a decarboxylase encoding gene in the *C. lactis* BGC, suggests that the peptide is a type B linaridin such as legonaridin or mononaridin (26, 50). In contrast however, the corynaridin gene cluster also misses a methyltransferase encoding gene, which makes it unique among the hitherto described peptides of the linaridin family. As N-terminal methylation was described to be essential for the antimicrobial activity of cypemycin (27) future studies need to address if methylation of corynaridin is catalyzed by other means.

Purified corynaridin was stable at temperatures up to 100°C but significantly lost activity after incubation with proteinase K or trypsin. Heat-stability is a favorable trait of many bacteriocins and was also shown for e.g. nisin (51). Interestingly, corynaridin has a bactericidal mode of action against *C. glutamicum* ATCC 13032 in liquid cultivations, but did not lead to the formation of pores as described for other class I bacteriocins such as nisin (52). As to our knowledge no receptor has been identified for the linaridins described so far, it remains to be investigated how these peptides exert their selective antimicrobial activity and in some cases cytotoxic activity against cancer cell lines (25). Characterization of corynaridin revealed antimicrobial activity against several other Actinobacteria comprising pathogenic and commensal *Corynebacterium* species as well as *M. luteus* and *C. acnes*, but only low or no activity against a selection of Firmicutes and Gram-negatives. *C. lactis* and other *Corynebacterium* species are frequently found in raw milk (13, 30) and some isolates of *Corynebacterium bovis* and *C. amycolatum* are causative agents of mastitis in dairy cows (53, 54). *C. lactis* was eventually linked to infections in companion animals, but is not considered a pathogen so far and the determinants of infection remain unclear (55, 56). Interestingly, a putative BGC similar to corynaridin exists in the genome of an isolate of *C. striatum* from an intensive care unit (49). *C. striatum* is a ubiquitous species and part of the human microbiome, but can also lead to infections in immunocompromised patients (57, 58). Moreover, Georgiou et al predicted linaridin BGCs in six *Corynebacterium* sp., including the ones in *C. lactis* and *C. striatum*, but also in a *C. diphtheriae* strain (7). These bacteria are not only phylogenetically related but might also be competitors in their respective habitats. Thus, secretion of bacteriocins by corynebacteria might be a hitherto largely neglected mechanism for intra-species competition as described for other bacterial groups, e.g. lactic acid bacteria, and might impact on the composition of (actino)bacterial communities in general (19, 59).

In conclusion, we identified a novel, heat stable bacteriocin produced by *C. lactis* RW3-42, which exerts narrow spectrum bactericidal activity against other Actinobacteria, mainly *Corynebacterium* species. Our experiments suggest that corynaridin has a yet undescribed, bactericidal mode of action that does not involve pore formation. As other linaridins also showed growth suppressing activity against cancer cell-lines, they might be promising candidates for biotechnological exploitation.

## 5 Declaration of competing interests

The authors declare no conflict of interest.

## 6 Acknowledgements

This study was partially funded by a grant of the German Ministry for Education and Research to CUR and BJE within the AMPLIFY consortium (Grant No. 031B0826A). The funding bodies had no role in the design of the study, analysis of the data, or writing of the manuscript.

## 7 Authors’ contribution

EP performed experimental work and was involved in writing, review and editing of the manuscript.

SJR performed the pHluorin experiments and analysis and was involved in writing, review and editing of the manuscript.

DW was involved in establishing the activity measurements and was involved in writing, review and editing of the manuscript.

BJE was involved in acquisition of funding, conceptualization of the study and writing, review and editing of the manuscript.

CUR was involved in acquisition of funding, conceptualization of the study and writing, review and editing of the manuscript.

OG performed conceptualization, experimental work, analyzed and visualized all data and writing, review and editing of the manuscript.

## Notes

### Competing Interest Statement

The authors have declared no competing interest.

## References

1. Alvarez-Sieiro P, Montalbán-López M, Mu D, Kuipers OP. 2016. Bacteriocins of lactic acid bacteria: extending the family. Appl Microbiol Biotechnol 100:2939–2951.

2. Chikindas ML, Weeks R, Drider D, Chistyakov VA, Dicks LM. 2018. Functions and emerging applications of bacteriocins. Curr Opin Biotechnol 49:23–28.

3. Cotter PD, Ross RP, Hill C. 2013. Bacteriocins-a viable alternative to antibiotics? Nat Rev Microbiol 11:95–105.

4. Field D, Ross RP, Hill C. 2018. Developing bacteriocins of lactic acid bacteria into next generation biopreservatives. Curr Opin Food Sci 20:1–6.

5. Soltani S, Hammami R, Cotter PD, Rebuffat S, Said L Ben, Gaudreau H, Bédard F, Biron E, Drider D, Fliss I. 2021. Bacteriocins as a new generation of antimicrobials: toxicity aspects and regulations. FEMS Microbiol Rev 45.

6. Riley MA, Wertz JE. 2002. Bacteriocins: Evolution, ecology, and application. Annu Rev Microbiol 56:117–137.

7. Georgiou MA, Dommaraju SR, Guo X, Mast DH, Mitchell DA. 2020. Bioinformatic and reactivity-based discovery of linaridins. ACS Chem Biol 15:2976–2985.

8. Goldbeck O, Weixler D, Eikmanns BJ, Riedel CU. 2021. In silico prediction and analysis of unusual lantibiotic resistance operons in the genus Corynebacterium. Microorganisms 9:1–18.

9. Gomes KM, Duarte RS, Bastos M do C de F. 2017. Lantibiotics produced by Actinobacteria and their potential applications (A review). Microbiol (United Kingdom) 163:109–121.

10. Poorinmohammad N, Bagheban-Shemirani R, Hamedi J. 2019. Genome mining for ribosomally synthesised and post-translationally modified peptides (RiPPs) reveals undiscovered bioactive potentials of actinobacteria. Antonie van Leeuwenhoek, Int J Gen Mol Microbiol 112:1477–1499.

11. Brennan NM, Brown R, Goodfellow M, Ward AC, Beresford TP, Simpson PJ, Fox PF, Cogan TM. 2001. Corynebacterium mooreparkense sp. nov. and Corynebacterium casei sp. nov., isolated from the surface of a smear-ripened cheese. Int J Syst Evol Microbiol 51:843–852.

12. Byrd AL, Belkaid Y, Segre JA. 2018. The human skin microbiome. Nat Rev Microbiol 16:143–155.

13. Hahne J, Kloster T, Rathmann S, Weber M, Lipski A. 2018. Isolation and characterization of Corynebacterium spp. from bulk tank raw cow’s milk of different dairy farms in Germany. PLoS One 13:1–16.

14. Bernard K. 2012. The genus Corynebacterium and other medically relevant coryneform-like bacteria. J Clin Microbiol 50:3152–3158.

15. Alibi S, Ferjani A, Boukadida J, Cano ME, Fernández-Martínez M, Martínez-Martínez L, Navas J. 2017. Occurrence of Corynebacterium striatum as an emerging antibiotic-resistant nosocomial pathogen in a Tunisian hospital. Sci Rep 7:1–8.

16. Cazanave C, Greenwood-Quaintance KE, Hanssen AD, Patel R. 2012. Corynebacterium prosthetic joint infection. J Clin Microbiol 50:1518–1523.

17. Otsuka Y, Ohkusu K, Kawamura Y, Baba S, Ezaki T, Kimura S. 2006. Emergence of multidrug-resistant Corynebacterium striatum as a nosocomial pathogen in long-term hospitalized patients with underlying diseases. Diagn Microbiol Infect Dis 54:109–114.

18. Silva-Santana G, Silva CMF, Olivella JGB, Silva IF, Fernandes LMO, Sued-Karam BR, Santos CS, Souza C, Mattos-Guaraldi AL. 2021. Worldwide survey of Corynebacterium striatum increasingly associated with human invasive infections, nosocomial outbreak, and antimicrobial multidrug-resistance, 1976–2020. Arch Microbiol 203:1863–1880.

19. Ramsey MM, Freire MO, Gabrilska RA, Rumbaugh KP, Lemon KP. 2016. Staphylococcus aureus Shifts toward commensalism in response to Corynebacterium species. Front Microbiol 7:1–15.

20. Abrehem K, Zamiri I. 1983. Production of a bacteriocin, ulceracin 378, by Corynebacterium ulcerans. Antimicrob Agents Chemother 24:262–267.

21. Pátek M, Hochmannová J, Nešvera J, Stránský J. Glutamicin CBII, a bacteriocin-like substance produced by Corynebacterium glutamicum. Antonie Van Leeuwenhoek 52:129–40.

22. Arnison PG, Bibb MJ, Bierbaum G, Bowers AA, Bugni TS, Bulaj G, Camarero JA, Campopiano DJ, Challis GL, Clardy J, Cotter PD, Craik DJ, Dawson M, Dittmann E, Donadio S, Dorrestein PC, Entian KD, Fischbach MA, Garavelli JS, Göransson U, Gruber CW, Haft DH, Hemscheidt TK, Hertweck C, Hill C, Horswill AR, Jaspars M, Kelly WL, Klinman JP, Kuipers OP, Link AJ, Liu W, Marahiel MA, Mitchell DA, Moll GN, Moore BS, Müller R, Nair SK, Nes IF, Norris GE, Olivera BM, Onaka H, Patchett ML, Piel J, Reaney MJT, Rebuffat S, Ross RP, Sahl HG, Schmidt EW, Selsted ME, Severinov K, Shen B, Sivonen K, Smith L, Stein T, Süssmuth RD, Tagg JR, Tang GL, Truman AW, Vederas JC, Walsh CT, Walton JD, Wenzel SC, Willey JM, Van Der Donk WA. 2013. Ribosomally synthesized and post-translationally modified peptide natural products: Overview and recommendations for a universal nomenclature. Nat Prod Rep 30:108–160.

23. Ma S, Zhang Q. 2020. Linaridin natural products. Nat Prod Rep 37:1152–1163.

24. Claesen J, Bibb MJ. 2011. Biosynthesis and regulation of grisemycin, a new member of the linaridin family of ribosomally synthesized peptides produced by Streptomyces griseus IFO 13350. J Bacteriol 193:2510–2516.

25. Minami Y, Yoshida K ichiro, Azuma R, Urakawa A, Kawauchi T, Otani T, Komiyama K, Omura S. 1994. Structure of cypemycin, a new peptide antibiotic. Tetrahedron Lett 35:8001–8004.

26. Rateb ME, Zhai Y, Ehrner E, Rath CM, Wang X, Tabudravu J, Ebel R, Bibb M, Kyeremeh K, Dorrestein PC, Hong K, Jaspars M, Deng H. 2015. Legonaridin, a new member of linaridin RiPP from a Ghanaian Streptomyces isolate. Org Biomol Chem 13:9585–9592.

27. Claesen J, Bibb M. 2010. Genome mining and genetic analysis of cypemycin biosynthesis reveal an unusual class of posttranslationally modified peptides. Proc Natl Acad Sci U S A 107:16297–16302.

28. Kanki K, Kazuhiko O, Toshiaki S, Kazuro S, Hong Y, Yoko T, Masahiko H, Toshio O. 1993. New Antibiotic, Cypemycin Taxonomy, Fermentation, Isolation and Biological Characteristics. J Antibiot (Tokyo) 46:1666–1671.

29. Shang Z, Winter JM, Kauffman CA, Yang I, Fenical W. 2019. Salinipeptins: Integrated Genomic and Chemical Approaches Reveal Unusual d -Amino Acid-Containing Ribosomally Synthesized and Post-Translationally Modified Peptides (RiPPs) from a Great Salt Lake Streptomyces sp. ACS Chem Biol 14:415–425.

30. Wiertz R, Schulz SC, Müller U, Kämpfer P, Lipski A. 2013. Corynebacterium frankenforstense sp. nov. and Corynebacterium lactis sp. nov., isolated from raw cow milk. Int J Syst Evol Microbiol 63:4495–4501.

31. Bachmann BJ. 1972. Pedigrees of some mutant strains of Escherichia coli K-12. Bacteriol Rev 36:525–557.

32. Bolotin A, Wincker P, Mauger S, Jaillon O, Malarme K, Weissenbach J, Ehrlich SD, Sorokin A. 2001. The complete genome sequence of the lactic acid bacterium Lactococcus lactis ssp. lactis IL1403. Genome Res 11:731–53.

33. Nilsen T, Nes IF, Holo H. 2003. Enterolysin A, a cell wall-degrading bacteriocin from Enterococcus faecalis LMG 2333. Appl Environ Microbiol 69:2975–84.

34. Glaser P, Frangeul L, Buchrieser C, Rusniok C, Amend A, Baquero F, Berche P, Bloecker H, Brandt P, Chakraborty T, Charbit A, Chetouani F, Couvé E, de Daruvar A, Dehoux P, Domann E, Domínguez-Bernal G, Duchaud E, Durant L, Dussurget O, Entian KD, Fsihi H, García-del Portillo F, Garrido P, Gautier L, Goebel W, Gómez-López N, Hain T, Hauf J, Jackson D, Jones LM, Kaerst U, Kreft J, Kuhn M, Kunst F, Kurapkat G, Madueno E, Maitournam A, Vicente JM, Ng E, Nedjari H, Nordsiek G, Novella S, de Pablos B, Pérez-Diaz JC, Purcell R, Remmel B, Rose M, Schlueter T, Simoes N, Tierrez A, Vázquez-Boland JA, Voss H, Wehland J, Cossart P. 2001. Comparative genomics of Listeria species. Science 294:849–52.

35. Cohn F. 1872. Untersuchungen über Bakterien. Beitr Biol Pflanz 1:127–224.

36. Rodríguez JM, Cintas LM, Casaus P, Martínez MI, Suárez A, Hernández PE. 1997. Detection of pediocin PA-1-producing pediococci by rapid molecular biology techniques. Food Microbiol 14:363–371.

37. Peters G, Locci R, Pulverer G. 1982. Adherence and growth of coagulase-negative staphylococci on surfaces of intravenous catheters. J Infect Dis 146:479–82.

38. Schäfer A, Tauch A, Jsger W, Kalinowski J, Thierbachb G, Piihler A. 1994. pK18mobsacB. Gene 145:49–5201.

39. Weixler D, Berghoff M, Ovchinnikov K V, Reich S, Goldbeck O, Seibold GM, Wittmann C, Bar NS, Eikmanns BJ, Diep DB, Riedel CU. 2022. Recombinant production of the lantibiotic nisin using Corynebacterium glutamicum in a two - step process. Microb Cell Fact 1–15.

40. Guzman LM, Belin D, Carson MJ, Beckwith J. 1995. Tight regulation, modulation, and high-level expression by vectors containing the arabinose PBAD promoter. J Bacteriol 177:4121–30.

41. van der Rest ME, Lange C, Molenaar D. 1999. A heat shock following electroporation induces highly efficient transformation of Corynebacterium glutamicum with xenogeneic plasmid DNA. Appl Microbiol Biotechnol 52:541–5.

42. Van Heel AJ, De Jong A, Song C, Viel JH, Kok J, Kuipers OP. 2018. BAGEL4: A user-friendly web server to thoroughly mine RiPPs and bacteriocins. Nucleic Acids Res 46:W278–W281.

43. Altschul S. 1997. Gapped BLAST and PSI-BLAST: a new generation of protein database search programs. Nucleic Acids Res 25:3389–3402.

44. Larkin MA, Blackshields G, Brown NP, Chenna R, Mcgettigan PA, McWilliam H, Valentin F, Wallace IM, Wilm A, Lopez R, Thompson JD, Gibson TJ, Higgins DG. 2007. Clustal W and Clustal X version 2.0. Bioinformatics 23:2947–2948.

45. Waterhouse AM, Procter JB, Martin DMA, Clamp M, Barton GJ. 2009. Jalview Version 2-A multiple sequence alignment editor and analysis workbench. Bioinformatics 25:1189–1191.

46. Haber M, Ilan M. 2014. Diversity and antibacterial activity of bacteria cultured from Mediterranean Axinella spp. sponges. J Appl Microbiol 116:519–532.

47. Crauwels P, Schäfer L, Weixler D, Bar NS, Diep DB, Riedel CU, Seibold GM. 2018. Intracellular phluorin as sensor for easy assessment of bacteriocin-induced membrane-damage in Listeria monocytogenes. Front Microbiol 9:1–10.

48. Mo T, Liu WQ, Ji W, Zhao J, Chen T, Ding W, Yu S, Zhang Q. 2017. Biosynthetic insights into linaridin natural products from genome mining and precursor peptide mutagenesis. ACS Chem Biol 12:1484–1488.

49. Roach DJ, Burton JN, Lee C, Stackhouse B, Butler-Wu SM, Cookson BT, Shendure J, Salipante SJ. 2015. A year of infection in the intensive care unit: prospective whole genome sequencing of bacterial clinical isolates reveals cryptic transmissions and novel microbiota. PLoS Genet 11:1–21.

50. Wang F, Wei W, Zhao J, Mo T, Wang X, Huang X, Ma S, Wang S, Deng Z, Ding W, Liang Y, Zhang Q. 2021. Genome mining and biosynthesis study of a type B linaridin reveals a highly versatile α-N-methyltransferase. CCS Chem 3:1049–1057.

51. Rollema HS, Kuipers OP, Both P, De Vos WM, Siezen RJ. 1995. Improvement of solubility and stability of the antimicrobial peptide nisin by protein engineering. Appl Environ Microbiol 61:2873–2878.

52. Bierbaum G, Sahl H-G. 2009. Lantibiotics: mode of action, biosynthesis and bioengineering. Curr Pharm Biotechnol 10:2–18.

53. Hommez J, Devriese LA, Vaneechoutte M, Riegel P, Butaye P, Haesebrouck F. 1999. Identification of nonlipophilic corynebacteria isolated from dairy cows with mastitis. J Clin Microbiol 37:954–957.

54. Watts JL, Lowery DE, Teel JF, Rossbach S. 2000. Identification of Corynebacterium bovis and other coryneforms isolated from bovine mammary glands. J Dairy Sci 83:2373–2379.

55. Antunes JMA de P, Ribeiro MG, Demoner L de C, Ramos JN, Baio PVP, Simpson-Louredo L, Santos CS, Hirata R, Ferioli RB, Romera ARC, Vieira VV, Mattos-Guaraldi AL. 2015. Cutaneous abscess caused by Corynebacterium lactis in a companion dog. Vet Microbiol 178:163–166.

56. Lim FS, Loong SK, Khoo JJ, Tan KK, Zainal N, Abdullah MF, Khor CS, AbuBakar S. 2018. Identification and characterization of Corynebacterium lactis isolated from Amblyomma testudinarium of Sus scrofa in Malaysia. Syst Appl Acarol 23:1838–1844.

57. Lee PP, Ferguson DA, Sarubbi FA. 2005. Corynebacterium striatum: An underappreciated community and nosocomial pathogen. J Infect 50:338–343.

58. McMullen AR, Anderson N, Wallace MA, Shupe A, Burnham CA. 2017. When good bugs go bad: Epidemiology and antimicrobial resistance profiles of Corynebacterium striatum, an emerging multidrug-resistant, opportunistic pathogen. Antimicrob Agents Chemother 61.

59. Kommineni S, Bretl DJ, Lam V, Chakraborty R, Hayward M, Simpson P, Cao Y, Bousounis P, Kristich CJ, Salzman NH. 2015. Bacteriocin production augments niche competition by enterococci in the mammalian gastrointestinal tract. Nature 526:719–722.

