## Supplementary data for "Identification, Characterization and Mode of Action of Corynaridin, a Novel Linaridin from *Corynebacterium lactis*"

### Corresponding authors.

E-mail:

Germany

**Table S1:** List of oligonucleotides used in this study.

| Primer | Sequence (5’ → 3’) | Purpose |
|---|---|---|
| pk19_fw<br>pk19_rev | GTATGTTGTGTGGAATTGTGAG<br>CAGGGTTTTCCCAGTCACGACG | Control primer flanking MCS pK19mobsacB |
| Do_crdA_fwd<br>Do_crdA_rev | CAAGCTTGCATGCCTGCAGGAAGAGGAACATTTGCCGC<br>GGTAGTCAGTTAGGTACTAACTAAAAATGATTGACG | Amplification of downstream region of *crdA* |
| Up_crdA_fwd<br>Up_crdA_rev | TTAGTACCTAACTGACTACCTTCCATTCTTAGG<br>TTGTAAAACGACGGCCAGTGGAGGAAGAATGCCGGAGAAAC | Amplification of upstream region of *crdA* |
| CorePeptide-HTHHyd-LinL_fwd | AGCGAATTCGAGCTCGGTACCCTAGATTCTAAGATAAGGAGG TAAATAAT<br>GCTCGCTACTGCAGTTAATTC | Contruction of pBAD33_crd |
| CorePeptide-HTHHyd-LinL_rev | TCAGTTCAGGCTCTTGCCGAGGAATGGG | Contruction of pBAD33_crd |
| ABC-SDR_fwd | TCGGCAAGAGCCTGAACTGAGAAAATTGAC | Contruction of pBAD33_crd |
| ABC-SDR_rev | CAAGCTTGCATGCCTGCAGGGGGAAATTACTCTGCTAAAAC | Contruction of pBAD33_crd |
| pBAD33_fw | ATTATTTGCACGGCGTCA | sequencing of *crd*-locus |
| pBAD33_rev | GTTTTATCAGACCGCTTCT | sequencing of *crd*-locus |
| S1 | GAGCTCGGTACCCTAGATTC | sequencing of *crd*-locus |
| S2 | GTGGCTGTCTAGGTACTAAC | sequencing of *crd*-locus |
| S3 | TCCACCAATCGGCTTAAAGG | sequencing of *crd*-locus |
| S4 | GAGGTCGTGTTCGGGTCTTG | sequencing of *crd*-locus |
| S5 | CGCAATGACAGCCTTTAACC | sequencing of *crd*-locus |
| S6 | GATTCAACCGAGAGCAGTTC | sequencing of *crd*-locus |
| S7 | GTATCGGTCGGGCAGTTGTG | sequencing of *crd*-locus |

**Table S2:** List of open reading frames contained in the *C. lactis* RW3-42 corynaridin gene cluster, the respective number of amino acids, putative TM-helices and homology to legonaridin biosynthesis proteins.

| orf[1] | amino acids | TM-helices | legonaridin homolog | amino acid identity (%) to legonaridin homologs |
|---|---|---|---|---|
| *crdA* | 69 | - | *legA* | 25 |
| *crdG* | 275 | 1 | *legH* | 32 |
| *crdE* | 324 | - | *legE* | 21 |
| *crdL* | 182 | - | *legF* | 26 |
| *crdT* | 550 | 6 | *legB* | 31 |
| *crdC* | 243 | - | *legC* | 25 |

[1]orf = open reading frame

```
LegA     1 MSVLAEFAN----TELVDV-EPGRLGSEATPTMITPLATLAT-------PEATPVGFAATS 49
CrdA     1 --MLATAVNSAQDRGLIPANYDGFLSMGSAPMTCTPTVTVRITVRIMRATKRTVRGAEP-- 57

LegA    50 ATAAAVNMITHDVTRH                                              65
CrdA    58 AASDRVETLVA---V-                                              69
```

**Figure S1:** CLUSTALW sequence alignment of LegA (Legonaridin) from *Streptomyces* sp. CT34 and CrdA (corynaridin) from *C. lactis* RW3-42.

```
LegH     1 VASLGTLVSALEMLSQHRKLADGELLSAQLESTRPEFAKR---FPRLTRALSSKK----  52
CrdG     1 ------MIDALKGL---QRVVQKP-LQA-----PLEFAERASAFSQLLATLEGISPKER  44

LegH    53 --AGV-------ALYGIQAGASAA--TMIW-----------------AHKRGVRAAGSA  83
CrdG    45 RFGGINDWDYTKYLFNSGGGISDAGVREIFVRCLATARIGASVVLLLPTGNNTRLVASS 103

LegH    84 VLAVTGAASRLRTPFGGDGADQLQQVINVVLAS-TGTFKDGDKGRDVAMRALALETTIS 141
CrdG   104 VSALSYLLGNRYTINGSDGAEQYSAI---ILASSALGRIDGGKNRDLAVDFIAAQTAFS 159

LegH   142 YVASGVVKLVSPVWLSGEAFSGVIRTHNYGDPNIYKLVHKYPMLGKLITWTTVAAEVGF 200
CrdG   160 YFVAGAVKSLGREWRNGTAVERVVRTEVYGNRIFYRFLRRNPRLSESLTYSTVVVEMLF 218

LegH   201 PLVFVLPKPAAKAYLGSMTLFHLGIGQFMGLNRFVLAFAATHPALLYVFDQSGRRPAVA 259
CrdG   219 PFLLLH-RGMRKVALASMFAFHAANVPLMGLGRFFIVFTSTYPAVMNSTNRLKGRFSD- 275

LegH   260 GNAVAALAPAAA                                                271
CrdG       ------------
```

**Figure S2:** CLUSTALW sequence alignment of LegH (HTTM-domain protein) from *Streptomyces* sp. CT34 and CrdG (HTTM-domain protein) from *C. lactis* RW3-42.

```
LegE     1 ------------------------------MRFTGKRERARRYLTHERPDGSLVEVDV-  28
CrdE     1 VTDFYRKTALITGGALGALKAGEMFGAFYADRFSRN------WPRHMRKTCARNEVAVY  53

LegE    29 --NIPPNARRMVLLDNGLGTTHEYWDWVCEALP--ADMGYVRFNRPGYGLSTPSKR--Y  81
CrdE    54 KRAGVDDSQGCILLVHGMGNSSVSLVRLGEEISELTGRTVIRYDRPGYGASRFCTDAPY 112

LegE    82 GLERHFALLQE----LRETYVADLPLVLAGHSLGGYFVAAYASLHPGAKEGVTGVVMID 136
CrdE   113 SVTQSVDELAEIIRWLHEQY---HWVTVVGHSFGGLLAY--LALMGLEEPSWCNALLIE 166

LegE   137 ATDVAHLRSSRRADI---------DRWSRQSMLME---Q-VFAVAGLSALRPALNQHKT 182
CrdE   167 PSH---LHEARRDPKRMLGVMLGVEELNRRHLLSPFGGELLDASIGPRALDGRRHPHVD 222

LegE   183 YRPEINRSYTAFLAQRRTWALAYRD-YRDAM-TYPELASVDSPLTVITAENNKGDNAAH 239
CrdE   223 AVRRERRSSRCVRATRREFTTLAKLLFDGTVIRAPQ---PSAEVHVVASARSVGDVRQK 278

LegE   240 QKVQAKLATLSKRSRQRCIDGSDHESLLSIQPHAAQVAEIIADEPPTGRADAERRKSAG 298
CrdE   279 ELFAEYI---CSPADMTILGETSHDTIVLDKDAVNRISQIVNG----GSIHAA------ 324

LegE   299 ARPAVKSAADREKEL                                             313
CrdE       ---------------
```

**Figure S3:** CLUSTALW sequence alignment of LegE (Alpha-beta hydrolase) from *Streptomyces* sp. CT34 and CrdE (Alpha-beta hydrolase) from *C. lactis* RW3-42.

```
LegF     1 VTVRVNTERLKTAAVTTSLAAWLVATAVAQMPEQ--RFDNLLKRGKLRIPTPNWRFFGP  57
CrdL     1 ---------MLRNVIEVVFGSWFTLTFLGQHPGGNHRSRGLLFRLSSTIFMPNWAFFAP  50

LegF    58 NPGVKDNHLLYRDVTDGKPGEWQEIPITRD---RAWYALAWNARNRSPKALFDAVQDVY 113
CrdL    51 NPGVYDDHLFYRVREGNEFSPWKEVVTSRNDDGPAWLSPFYSGASRRSKGIIDIFSTLE 109

LegF   114 VRSAAYGTAMEPVVQSSGYQLLSGYIRHHLPHAEGASHSQFLVMYSYLAAPEARQIEPI 172
CrdL   110 SLASPTLSRTDCNVIQAGRQAIANFMVVNADEI-SPHTSEYEVMLVR-AAGYAQDEDPV 166

LegF   173 FVSREFPLEDEGVVQPEPYPVAA                                     195
CrdL   167 FYYR-FAVRNGIADAPH------                                     182
```

**Figure S4:** CLUSTALW sequence alignment of LegF (LinL protein) from *Streptomyces* sp. CT34 and CrdL (LinL protein) from *C. lactis* RW3-42.

```
LegB     1 -----------MLLITGVLLGLLGVGASLTQPWAIGKLIEAAGKGESLAFPIVRLVGLF  48
CrdT     1 MTAFNLLRNNAGLITLGILLSLLSTAVTLFQPALVGQLISGVSSGELQQ-PLMLLLLTV  58

LegB    49 CLGAVFSALQAYVIGRAGENIVYDVRQVLTGRLLRADLTEFGKRPQGDILTRTVTDTSL 107
CrdT    59 LGTSVLTAATMYVVSIAADRTVRDMRKKITNHLLYLRVRELEKGGSGSFTTRVTSDTSI 117

LegB   108 VKIALSQSLAQLIVSGATVIGGVVLMFLIDVRLMLITMGCLGVASLLSLSIARKLRRVA 166
CrdT   118 VSTAFSSTLTDFVGGFTVIIGALIYMAVVDWKLLLVVIAVLLVALAIIVSISSSLQNLS 176

LegB   167 VQNRDDTGEFGTAVQRVLAALPTVKASRAETRETARIGRLAERARRSGIRVNAYNALFM 225
CrdT   177 SKVQDHLAALGEILQSALSAIRTIKAFRVETKVIGNLSAEIDHAYRNRRRMSFVEAVLE 235

LegB   226 PSMNVGTQGALAVVVGAGMAWVARGEMNMADLTTFIMYLFQLVSPLVTFFMGVGQFIQG 284
CrdT   236 PLSTVASYMALLAVVLFGSIRLSNGDLSGEALTVFVTALFLMLAPIVQVSQSLGAFFEA 294

LegB   285 RAAIQRVDELAQMPQEGNGAQAARSSVPIGPLPQHLPAVEFHDVRFGYRRGEEGERTQV 343
CrdT   295 RGALDRINQLLRLEVEDSTE--SSSSLCTGPT--PLGTIEFDAVSYVR------EKRTI 343

LegB   344 LHGISLTVPARGLTAVVGPSGAGKTSMFQLIERFYALDGGVILLGGQNIGTLPLDTVRG 402
CrdT   344 LDNATVTVKSGEKVALTGASGSGKTSILSLLLKFYDVSAGHIRIGGKDLEDWNRRDLRA 402

LegB   403 LVGYVQQDSATMRGTVRENLTYAHPHASEDDIWEA-VEMAGLTSVVANLPKGLDTELGD 460
CrdT   403 MVTYVEQEPDLLSGTLRENLILGTGESFDDETLIAMLGQFGLE--NFASVDGLSRSVGS 459

LegB   461 QGSGLSGGQRQRLCIARTLLQKPAVMLLDEATSNLDSDSERDFRDVLRRVSQQCAVIAI 519
CrdT   460 GNSGLSGGERQRVAIIRAVLQNAPVILVDEPTSALDSASAELSMSRL--LETDSTVIFT 516

LegB   520 AHRISTVVDAEKIVVLEGGRVRATGVHRELMEHDELYRRLAGSQLHAGAAPVATATHWV 578
CrdT   517 SHDANIVNLAERTLVVVDGRIVENSPNSRKSSNA------------------------- 550

LegB   579 SGEPGAGPLASALGWHQWGSGLNEQTVRLRPIRIEEQW                      616
CrdT       --------------------------------------
```

**Figure S5:** CLUSTALW sequence alignment of LegB (ABC transporter) from *Streptomyces* sp. CT34 and CrdT (ABC transporter) from *C. lactis* RW3-42.

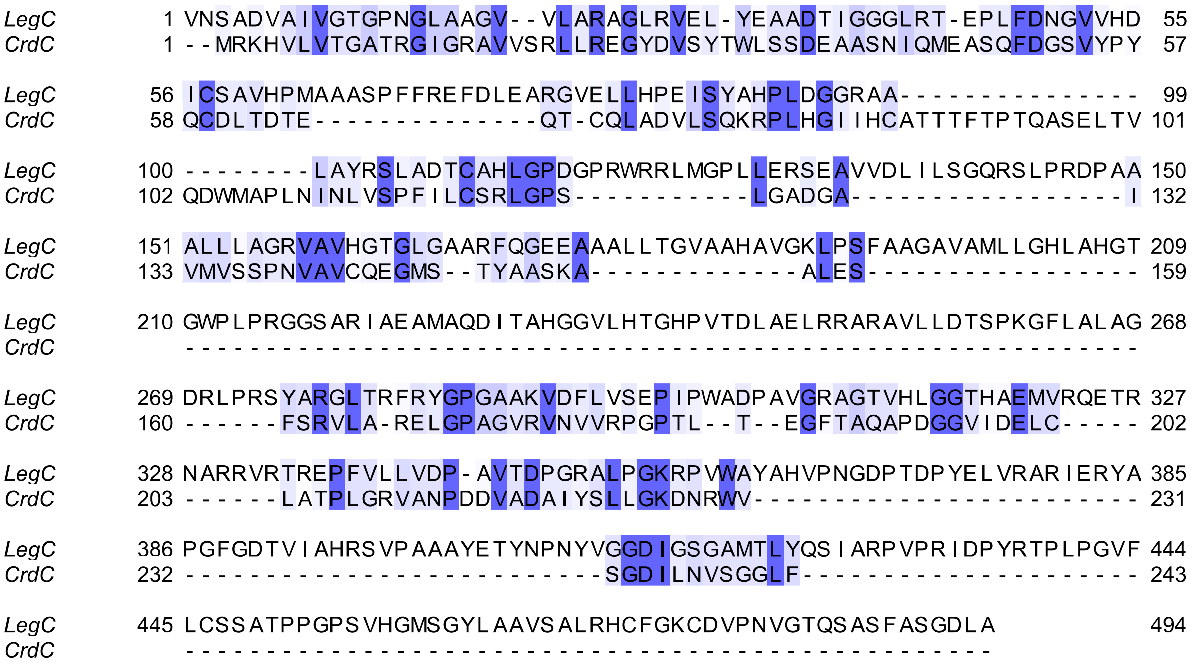


**Figure S6:** CLUSTALW sequence alignment of LegC (SDR oxidoreductase) from *Streptomyces* sp. CT34 and CrdC (SDR oxidoreductase) from *C. lactis* RW3-42.

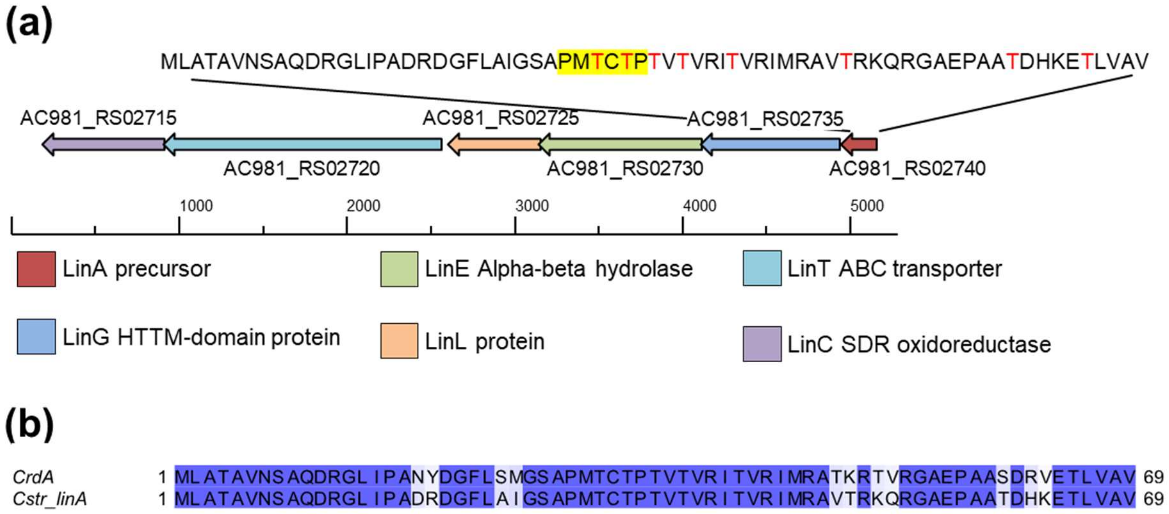


**Figure S7:** Putative linaridin gene cluster in *C. striatum* 1329_CAUR. (a) Gene cluster and annotation according to BLASTP analyses. (b) Sequence alignment of the *C. striatum* precursor peptide AC981_RS02740 (Cstr_linA) and corynaridin (CrdA) of *C. lactis*.

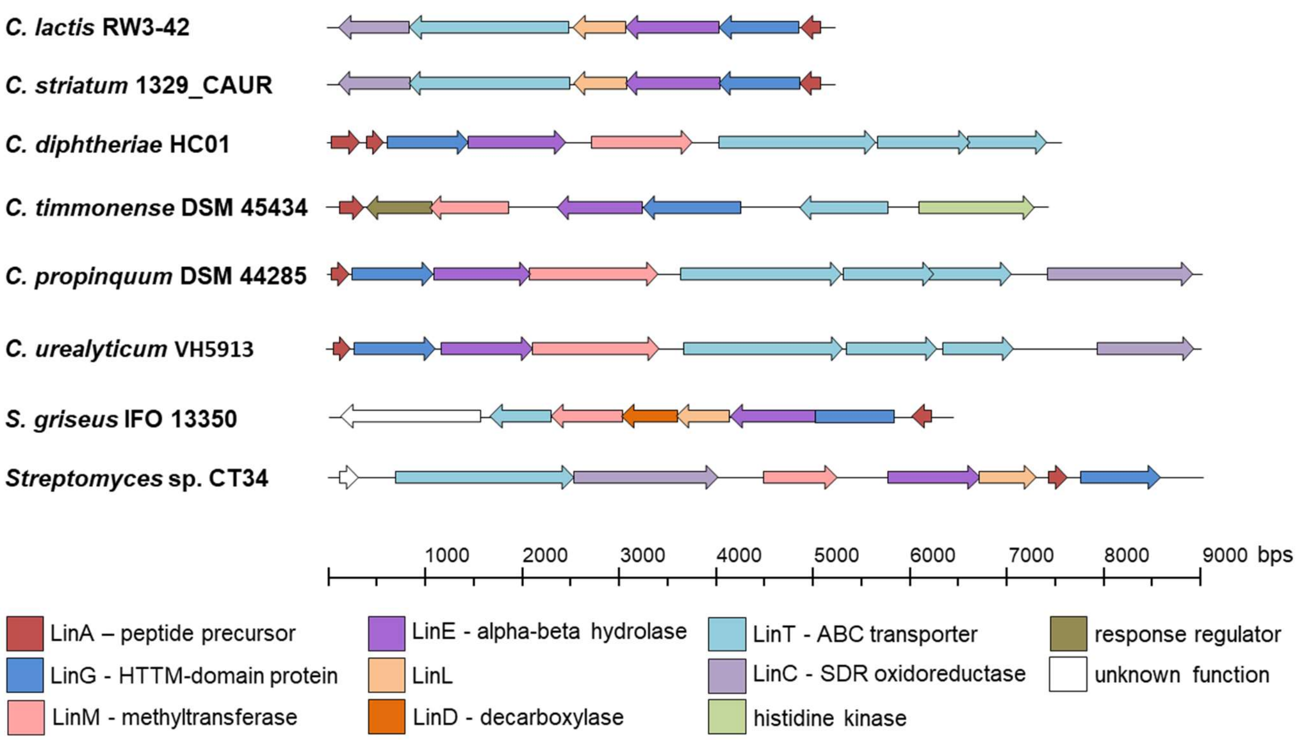


**Figure S8:** Putative linaridin gene clusters in *Corynebacterium* strains (1) compared with BGCs for other bacteriocins of the linaridin family. Gene cluster and annotation according to BLASTP analyses.

| | |
|---|---|
| ***C. lactis*** **RW3-42** | MLATAVNSAQDRGLIPANYDGFLSMGSAPMTCTPTVTVRITVRIMRATKRTVRGAEPAASDRVETLVAV |
| ***C. striatum*** **1329_CAUR** | MLATAVNSAQDRGLIPADRDGFLAIGSAPMTCTPTVTVRITVRIMRAVTRKQRGAEPAATDHKETLVAV |
| ***C. diphtheriae*** **HC01** | MQPFSGFSLSSDERLAYFSIGHFILFQTTFQGVDLSALSQLVNDNTAFRGRLDGMAETALPPVTATPAIAVTAAKAAGAVIGGGAVSGAAYAAYRAVIK |
| | MSRLSTLVNNDQITLEQVHSPVMATPAAFGAGVVAGAKACGALVAAAGVGAAIASATKK |
| ***C. timmonense*** **DSM 45434** | MRNNITKFADEADLQQVGSLYGDSEPVLATIITTTITATTATTATTAAGYHSSRPDEVMAGIDEDAPVSEMLAARKDAMLV |
| ***C. propinquum*** **DSM 44285** | MSRLSTLVNNDQITLEQVHSPVMATPAAFGAGVVAGAKACGALVAAAGVGAAIATATKE |
| ***C. urealyticum*** **VH5913** | MSRLSTLVNNDQITLEQVHSPVMATPAAFGAGVVAGAKACGALVAAAGVGAAIASATKK |
| ***S. griseus*** **IFO 13350** | MRSEMTLTSTNSAEALAAQDFANTVLSAAAPGFHADCETPAMATPATPTVAQFVIQGSTICLVC |
| ***Streptomyces*** **sp. CT34** | MSVLAEFANTELVDVEPGRLGSEATPTMITPLATLATPEATPVGFAATSATAAAVNMITHDVTRH |

**Figure S9:** Predicted precursor peptide sequences in *Corynebacterium* strains (1) compared with BGCs for other bacteriocins of the linaridin family. Threonine residues are displayed in red, serine residues in blue letters. The predicted hexapeptide cleavage site PxxxTP is highlighted in yellow.

**Table S3:** Results of cross-streak and spot-on-lawn assays with different indicator bacteria. Cross streak assay with *C. lactis* RW3-42 streaked in the middle of a BHI agar plate and indicator bacteria lateral to it. Nisin (250 µg/mL) and corynaridin RPC-fraction (>300 µg/mL) were spotted in 10 µl drops onto the plates. Spots were documented using an iBright imaging device or a light-table and camera. n.d. = not detected.

| Strain | Cross streak | Nisin | Corynaridin |
|---|---|---|---|
| *Bacillus subtilis* DSM 402 | 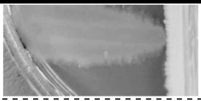 | 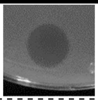 | n.d. |
| *Corynebacterium ammoniagenes* DSM 20306 | 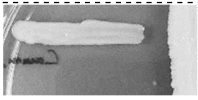 | 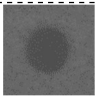 | 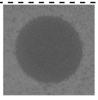 |
| *Corynebacterium amycolatum* DSM 6922 | 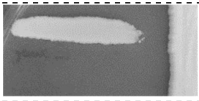 | 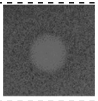 | 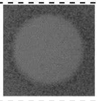 |
| *Corynebacterium canis* DSM 45402 | 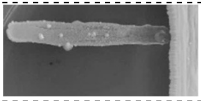 | n.d. | n.d. |
| *Corynebacterium casei* DSM 44701 | 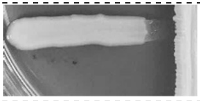 | 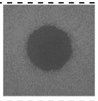 | 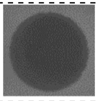 |
| *Corynebacterium efficiens* DSM 44549 | 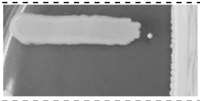 | 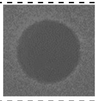 | 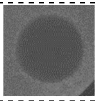 |
| *Corynebacterium glutamicum* ATCC 13032 | 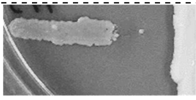 | 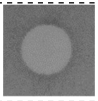 | 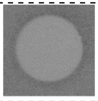 |
| *Corynebacterium lipophiloflavum* DSM 44291 | 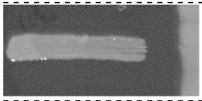 | 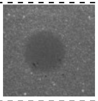 | 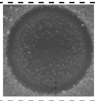 |
| *Corynebacterium striatum* DSM 20668 | 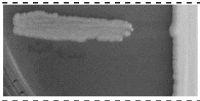 | 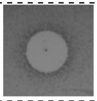 | 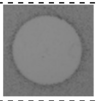 |
| *Corynebacterium xerosis* DSM 20743 | 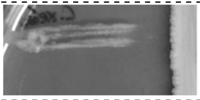 | 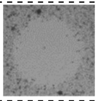 | 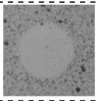 |
| *Cutibacterium acnes* DSM 16379 | 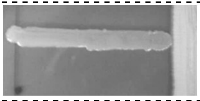 | 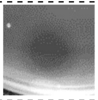 | 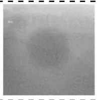 |
| *Escherichia coli* K12 MG1655 | 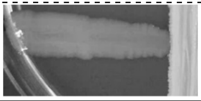 | n.d. | n.d. |

➔ continued

**continuation of table S3:**

| strain | cross streak | nisin | corynaridin |
|---|---|---|---|
| *Lactobacillus plantarum* DSM 1055 | 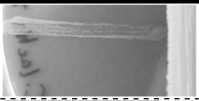 | 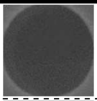 | n.d. |
| *Lactococcus lactis* IL1403 | 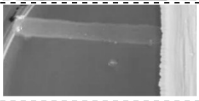 | 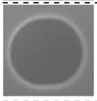 | 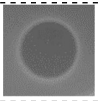 |
| *Listeria innocua* LMG2785 | 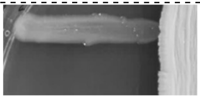 | 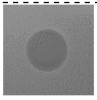 | 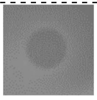 |
| *Listeria monocytogenes* EGD-e | 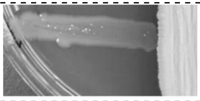 | 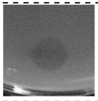 | n.d. |
| *Micrococcus luteus* DSM 20030 | 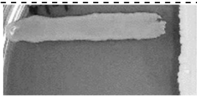 | 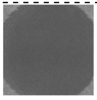 | 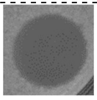 |
| *Pediococcus acidilactici* 347 | 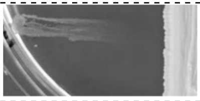 | 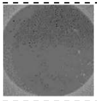 | n.d. |
| *Pseudomonas fluorescens* DSM 50090 | 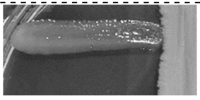 | n.d. | n.d. |
| *Staphylococcus aureus* ATCC 29213 | 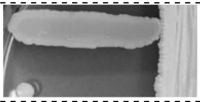 | 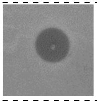 | n.d. |
| *Staphylococcus epidermidis* DSM 3269 | 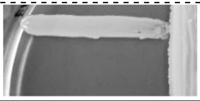 | 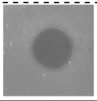 | n.d. |

#### References


1. Georgiou MA, Dommaraju SR, Guo X, Mast DH, Mitchell DA. 2020. Bioinformatic and reactivity-based discovery of linaridins. ACS Chem Biol 15:2976–2985.
